# Grant writing coaching groups: A randomized trial to discern the influences of coaching duration and engagement of scientific mentors on participants’ proposal submission and success rates

**DOI:** 10.64898/2026.07.29.741400

**Authors:** Kolawole S Okuyemi, Anne Marie Weber-Main, Melanie Steiner, Jeffrey Engler, Harlan P Jones, Wenxian Zhou, Patrick O Monahan, Andrew K Langi, Richard McGee

## Abstract

**Introduction:** The acquisition of National Institutes of Health (NIH) grants is an increasingly competitive undertaking. Many early-career biomedical investigators lack sufficient grant writing skills. This pragmatic, 2x2 factorial, group-randomized trial tested the effects of coaching duration and mode of engagement of participants’ scientific mentors in a grant writing coaching group intervention on grant application outcomes.

**Methods:** A national US sample of 367 faculty and postdoctoral fellows, organized into 76 coaching groups across 6 cohorts (2020-2022), was randomized to 1 of 4 arms that differed by coaching duration (5 months vs 5 months plus 18 months of access to 1:1 coaching) and mode of scientific mentor engagement (structured vs unstructured). Primary outcomes were national-level grant application submissions and awards, and success rate among submitters, assessed via surveys and verified through award databases 30 months after study initiation.

**Results:** Among 271 faculty and 96 postdoctoral fellows, 69% submitted at least 1 national application and 32% were awarded at least 1 grant. Submission rates did not differ by experimental arm. Among faculty, the structured engagement of a mentor in coaching was associated with significantly higher odds of any award (adjusted OR = 1.87) and any NIH mentored career development (K) award (adjusted OR = 3.32), and with higher success rates for these proposal types. Extended coaching duration showed a positive but not statistically significant association with faculty awards after adjustment. Neither factor significantly affected outcomes for postdoctoral fellows. Faculty success rates exceeded national NIH-reported success rates for research project grants equivalent to the R01 mechanism (43% vs 17.6%) and K awards (51% vs 36.5%).

**Conclusion:** Participation in a national, cross-institutional, group coaching model yielded high grant submission outcomes, and award rates were higher relative to NIH benchmarks. Structured integration of scientific mentors with the coaching intervention had a greater effect than extended coaching duration on faculty outcomes.

## Introduction

Sustaining a vibrant research enterprise to advance human health depends on a highly trained biomedical workforce to conduct investigator-initiated, grant-funded research. However, securing grant funding is an increasingly competitive undertaking. Funding success rates at the National Institutes of Health (NIH) declined from 25% in 1998 to 17% in 2024 among new R01 and R01-equivalent awards amidst a doubling of investigators who apply [1]. Success rates for NIH research career development awards similarly declined from 50% to 31% over the same period [2]. Many early-stage investigators enter their faculty roles with strong disciplinary expertise but without sufficient grant writing skills. Opportunities to attain these skills can be highly variable, dependent on institutional research infrastructures and on the availability and engagement of experienced mentors.

To help meet this training need, the NIH and other extramural agencies have invested in professional development programs at academic institutions, centers, and consortia across the US [3–8]. From 2014 to 2019, the NIH-funded National Research Mentoring Network (NRMN) implemented 6 distinct national models of grant writing coaching programs. In each model, participants engaged in several months of sustained, intensive coaching with experienced investigators, nested within a small cohort of peers who were actively writing grant applications [9–11]. The models differed in features such as cohort size, participant eligibility, approaches to giving feedback, and coaching group duration. Our evaluation of these models affirmed the overall value of group coaching to support proposal development and grant acquisition, while identifying strengths and weaknesses of each variation [10].

We applied these insights to design an enhanced group coaching intervention that incorporated some of the previous models’ strongest elements (eg, consideration of participants’ writing readiness and time commitments, coaches with success in securing grant funding, alignment of group members and coach by research area, focus on developing the entire application rather than a few sections) [6]. Additionally, we observed that individuals who participated in groups without the involvement of a scientifically-aligned mentor often struggled, and that a subset of participants needed a more extended period of coaching to develop their application. We hypothesized that 2 features that had not yet been empirically tested—coaching duration and involvement of a scientific mentor—could be leveraged to increase participants’ grant writing success.

We therefore designed a pragmatic, 2x2 factorial, group-randomized trial to answer 2 research questions aimed at optimizing our prototype intervention’s effectiveness: First, what duration, or dose, of the intervention is most effective? Second, is there a benefit to a participant’s scientific mentor being directly integrated into the group coaching process? We hypothesized that grant application outcomes, defined as submissions and awards, would be greater for participants who had 18 months of extended access to a skilled grant writing coach in addition to the regular dose of 5 months of group coaching, and for participants whose scientific mentor had direct, or structured, engagement with the coaching process versus unstructured, or no direct, interaction with the coaching process. We report trial outcomes here, focusing on application submissions to and grant awards from national funding sources.

## Methods

The study received an exemption determination by the Institutional Review Board (IRB) at the University of Utah (#00113440), and the IRB waived documentation of signed consent. The consent process was administered electronically using Research Electronic Data Capture (REDCap), a secure web-based data capture platform [12]. We previously published the complete study protocol [6] and offer an abbreviated description here.

### Study design, participant eligibility, and group randomization

We recruited a national US-based sample of early-career investigators (faculty and postdoctoral fellows) between October 1, 2019, and June 30, 2022, to participate in a pragmatic, unblinded, group-randomized trial (2x2 factorial design) to test variations of a group coaching intervention for the development of NIH and comparable national grant applications. Eligible for inclusion were individuals without prior major independent research funding (eg, NIH R01) who were actively writing career development or research (K- or R-type) applications; had the necessary research background, preliminary data, and institutional resources to realistically compete for federal or comparable national funding; and had adequate time to commit to study participation. We recruited 6 distinct cohorts, staggered to begin the trial in either January or June of 2020, 2021, and 2022.

Participants enrolled as dyads with a self-selected, scientifically aligned mentor, hereafter called scientific advisor (SA). An SA was defined as an individual with relevant content expertise who agreed to provide scientific feedback on the participant’s developing proposal as a complement to the intervention’s group coaching process. Dyads were placed into coaching groups (4 to 5 dyads each) based on broad similarities in research type. Coaching groups were randomized to 1 of 4 study arms that differed on 2 factors: duration of coaching support (regular versus extended dose) and mode of engaging SAs with the group coaching process (structured versus unstructured). Cohorts began with an equal number of groups in each study arm.

### Interventions

The regular dose of coaching experienced by all participants consisted of 8 biweekly virtual group coaching sessions held over 5 months of active proposal writing. Coaches were NIH-funded established investigators who had received training in the intervention’s coaching approaches [6]. At each session, participants received written and/or oral feedback on their drafts from their coach and other group members. If at least 1 participant within a coaching group completed an application, the group held an NIH-style mock study section that engaged reviewers identified by the participants or coach [6]. In the extended dose arms, participants had access to one-on-one coaching (up to 10 hours) after completion of regular group coaching. At the start of the extended dose period, coaches checked in with each group member to assess their anticipated needs for ongoing support. Thereafter, participants reached out to their coaches as needed.

To test modes of SA engagement, SAs in the structured engagement arms agreed to 3 touchpoints with the group coaching intervention: (1) provide the coach with a written review of an early draft of the participant’s Specific Aims page; (2) attend at least 1 group coaching session; and (3) participate in 1 virtual meeting with the coach and participant approximately midway through the intervention to assess writing progress and compare feedback. SAs were also invited to attend the group’s NIH-style mock study section, if held. In the unstructured engagement arms, SAs and coaches were not known to each other and interacted independently with the participant.

### Data collection

Self-reported grant application submission and award data were captured via online REDCap surveys administered at 6, 12, 18, and 24 months after the initiation of coaching for cohorts 1 through 6. Surveys remained open for 6 weeks; participants received multiple reminder emails. Outcome survey response rates were 81% (6 months), 68% (12 months), 65% (18 months), and 64% (24 months). Award data for participants who reported an application to a federal agency or national foundation were independently confirmed using public databases, such as NIH RePORTER [13] and foundation websites, for 30 months after initiation of coaching. Data collection began in June 2020 and was completed in October 2024; cohort 6 awards were verified through December 2024.

### Outcome variables

The 3 primary outcomes, measured at the participant level for all participants, faculty only, and postdoctoral fellows only, were as follows: (1) percentages of participants who submitted at least 1 national-level application within 24 months of coaching initiation (yes/no); (2) percentages of participants who were awarded at least 1 national-level grant within 30 months of coaching initiation (yes/no); and (3) success rates for national-level applications, defined as the percentage of participants who were funded among the subset who submitted. We also analyzed, as secondary outcomes, the submission, award, and success rates for specific categories of grants (any NIH K, NIH R01, NIH and NSF R01-like, Other NIH Research, Other Federal Research, and National Foundation; S1 Table).

### Statistical analyses

All tests were 2-sided. The 4 study arms were assessed for imbalance on baseline characteristics using ANOVA F test and Wilcoxon Rank Sum Test for continuous characteristics, and Pearson chi-square tests (or Freeman-Halton extension of Fisher’s exact test when > 20% of cells had expected counts < 5) for categorical characteristics.

Bivariable (unadjusted) and multivariable (adjusted) logistic regression analyses were used to test the 2 experimental factors (regular versus extended dose, structured versus unstructured SA engagement). Covariates were adjusted for in multivariable models if they were statistically (*P* < .05) imbalanced at baseline (age, gender, research focus, any prior training grant submission, any prior research grant submission) or theoretically important (underrepresented in medicine [URM] versus non-URM, number of prior peer-reviewed publications, and career level in the all participant and faculty models). We tested 2-way interaction effects in multivariable models between the 2 experimental factors, and between each experimental factor and each baseline characteristic (listed above, each was pre-specified to be a hypothesized potential intervention moderator); if significant, the SLICE option in SAS LOGISTIC provided simple effects (ie, experimental effects separately for levels of categorical moderators or for specified values for continuous moderators). These subgroup analyses were examined only if the interaction test was significant.

## Results

Across the 6 cohorts, 713 individuals applied to join the study, of whom 525 were deemed eligible. Among individuals meeting eligibility criteria, 158 were not enrolled because the cohorts had reached capacity. Ultimately, 367 individuals were enrolled in the study and organized into 76 coaching groups (Fig 1).

**Fig 1.**
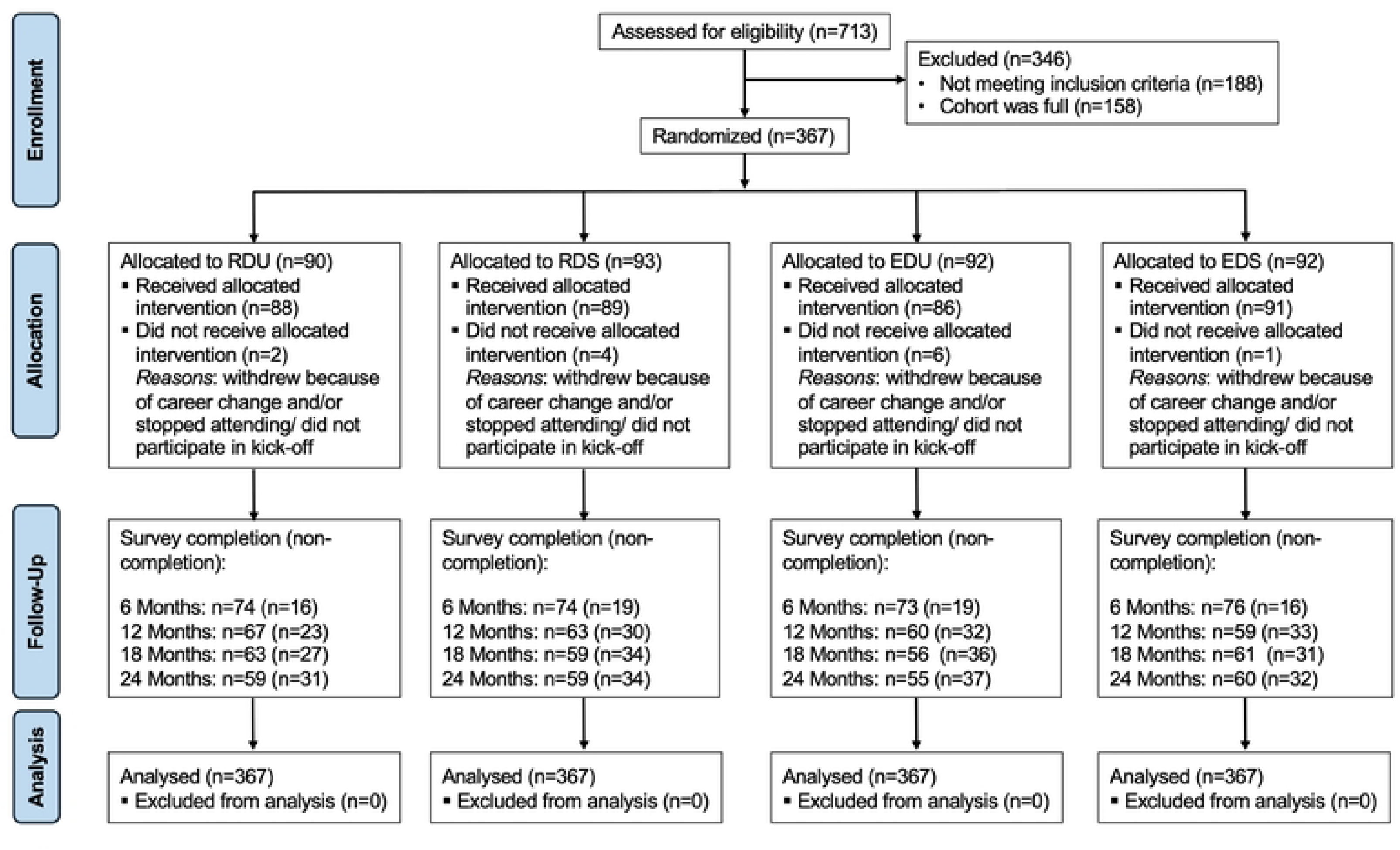
CONSORT Diagram Depicting Participant Flow Through the Trial. RDU = regular dose unstructured; RDS = regular dose unstructured; EDU = extended dose unstructured; EDS = extended dose structure.

### Sample characteristics

Table 1 shows the demographic and research characteristics of participants overall and by study arm. Variables with statistically significant differences among arms (except for prior training awards due to collinearity with prior training submissions) were included in multivariate models, along with theoretically important covariates.

**Table 1.**
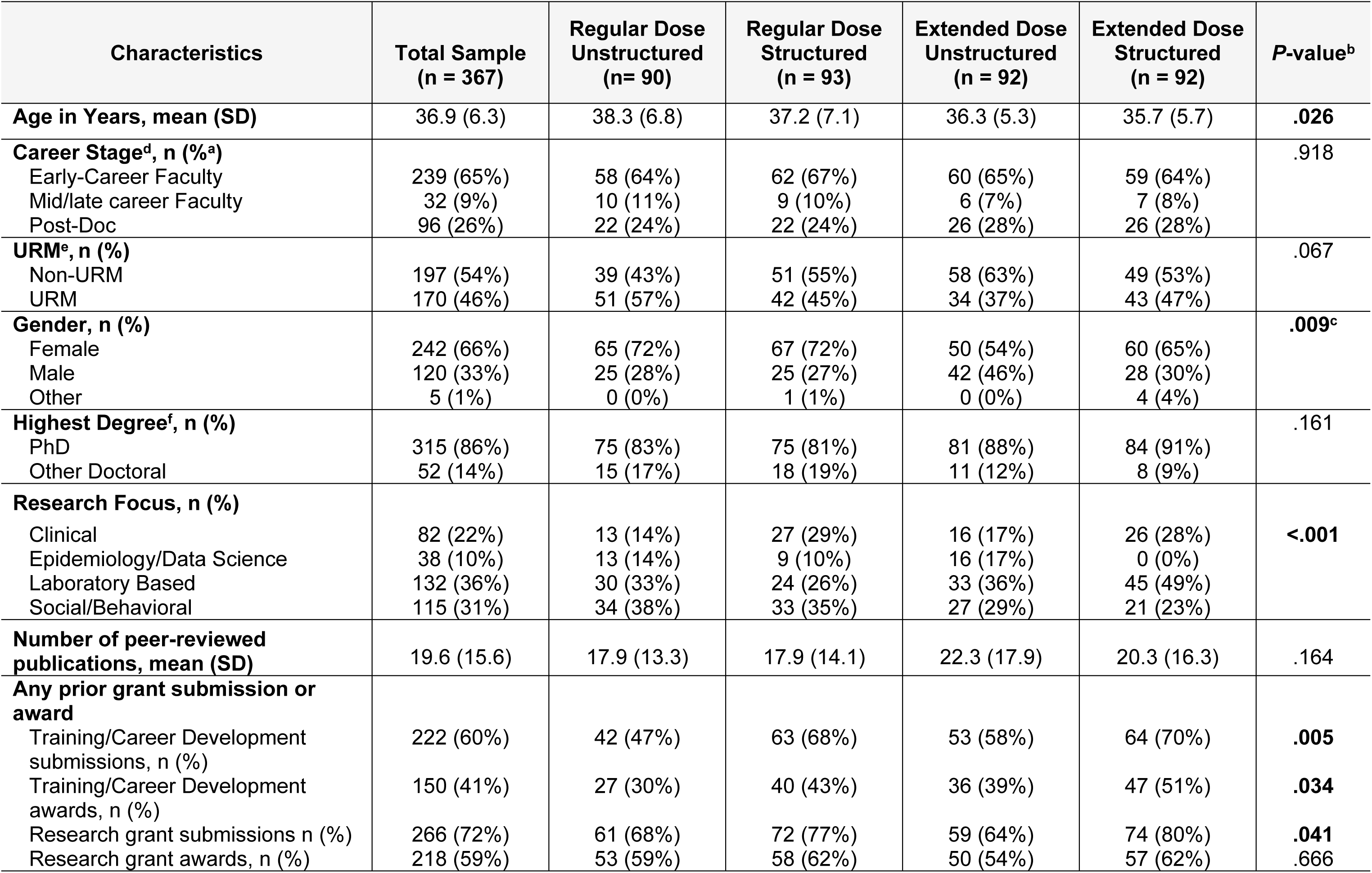
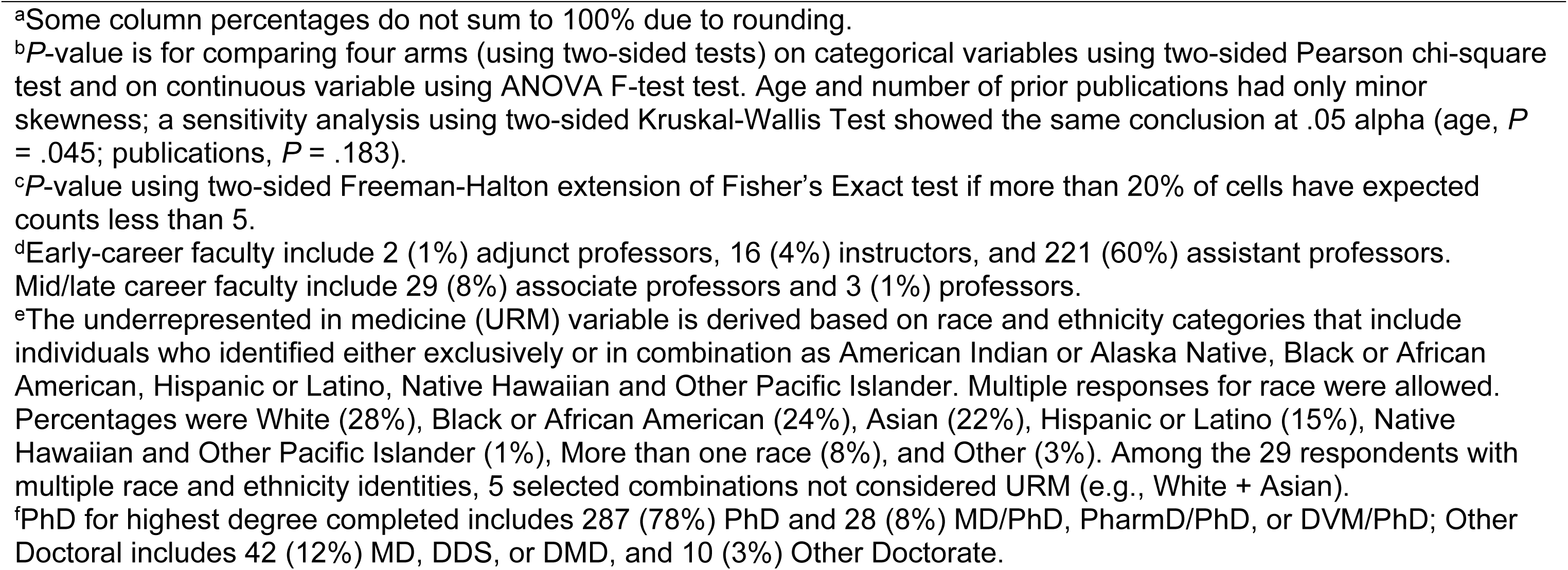
Characteristics of 367 US-Based Early-Career Faculty and Postdoctoral Fellows Enrolled in 2000-2022 Grant Coaching Intervention Study.

| Characteristics | Total Sample<br>(n = 367) | Regular Dose<br>Unstructured<br>(n= 90) | Regular Dose<br>Structured<br>(n = 93) | Extended Dose<br>Unstructured<br>(n = 92) | Extended Dose<br>Structured<br>(n = 92) | P-value<sup>b</sup> |
| --- | --- | --- | --- | --- | --- | --- |
| **Age in Years, mean (SD)** | 36.9 (6.3) | 38.3 (6.8) | 37.2 (7.1) | 36.3 (5.3) | 35.7 (5.7) | **.026** |
| **Career Stage<sup>d</sup>, n (%<sup>a</sup>)** |  |  |  |  |  | .918 |
| Early-Career Faculty | 239 (65%) | 58 (64%) | 62 (67%) | 60 (65%) | 59 (64%) |  |
| Mid/late career Faculty | 32 (9%) | 10 (11%) | 9 (10%) | 6 (7%) | 7 (8%) |  |
| Post-Doc | 96 (26%) | 22 (24%) | 22 (24%) | 26 (28%) | 26 (28%) |  |
| **URM<sup>e</sup>, n (%)** |  |  |  |  |  | .067 |
| Non-URM | 197 (54%) | 39 (43%) | 51 (55%) | 58 (63%) | 49 (53%) |  |
| URM | 170 (46%) | 51 (57%) | 42 (45%) | 34 (37%) | 43 (47%) |  |
| **Gender, n (%)** |  |  |  |  |  | **.009<sup>c</sup>** |
| Female | 242 (66%) | 65 (72%) | 67 (72%) | 50 (54%) | 60 (65%) |  |
| Male | 120 (33%) | 25 (28%) | 25 (27%) | 42 (46%) | 28 (30%) |  |
| Other | 5 (1%) | 0 (0%) | 1 (1%) | 0 (0%) | 4 (4%) |  |
| **Highest Degree<sup>f</sup>, n (%)** |  |  |  |  |  | .161 |
| PhD | 315 (86%) | 75 (83%) | 75 (81%) | 81 (88%) | 84 (91%) |  |
| Other Doctoral | 52 (14%) | 15 (17%) | 18 (19%) | 11 (12%) | 8 (9%) |  |
| **Research Focus, n (%)** |  |  |  |  |  | **<.001** |
| Clinical | 82 (22%) | 13 (14%) | 27 (29%) | 16 (17%) | 26 (28%) |  |
| Epidemiology/Data Science | 38 (10%) | 13 (14%) | 9 (10%) | 16 (17%) | 0 (0%) |  |
| Laboratory Based | 132 (36%) | 30 (33%) | 24 (26%) | 33 (36%) | 45 (49%) |  |
| Social/Behavioral | 115 (31%) | 34 (38%) | 33 (35%) | 27 (29%) | 21 (23%) |  |
| **Number of peer-reviewed publications, mean (SD)** | 19.6 (15.6) | 17.9 (13.3) | 17.9 (14.1) | 22.3 (17.9) | 20.3 (16.3) | .164 |
| **Any prior grant submission or award** |  |  |  |  |  |  |
| Training/Career Development submissions, n (%) | 222 (60%) | 42 (47%) | 63 (68%) | 53 (58%) | 64 (70%) | **.005** |
| Training/Career Development awards, n (%) | 150 (41%) | 27 (30%) | 40 (43%) | 36 (39%) | 47 (51%) | **.034** |
| Research grant submissions n (%) | 266 (72%) | 61 (68%) | 72 (77%) | 59 (64%) | 74 (80%) | **.041** |
| Research grant awards, n (%) | 218 (59%) | 53 (59%) | 58 (62%) | 50 (54%) | 57 (62%) | .666 |

<sup>a</sup>Some column percentages do not sum to 100% due to rounding.
<sup>b</sup>*P*-value is for comparing four arms (using two-sided tests) on categorical variables using two-sided Pearson chi-square test and on continuous variable using ANOVA F-test test. Age and number of prior publications had only minor skewness; a sensitivity analysis using two-sided Kruskal-Wallis Test showed the same conclusion at .05 alpha (age, *P* = .045; publications, *P* = .183).
<sup>c</sup>*P*-value using two-sided Freeman-Halton extension of Fisher's Exact test if more than 20% of cells have expected counts less than 5.
<sup>d</sup>Early-career faculty include 2 (1%) adjunct professors, 16 (4%) instructors, and 221 (60%) assistant professors. Mid/late career faculty include 29 (8%) associate professors and 3 (1%) professors.
<sup>e</sup>The underrepresented in medicine (URM) variable is derived based on race and ethnicity categories that include individuals who identified either exclusively or in combination as American Indian or Alaska Native, Black or African American, Hispanic or Latino, Native Hawaiian and Other Pacific Islander. Multiple responses for race were allowed. Percentages were White (28%), Black or African American (24%), Asian (22%), Hispanic or Latino (15%), Native Hawaiian and Other Pacific Islander (1%), More than one race (8%), and Other (3%). Among the 29 respondents with multiple race and ethnicity identities, 5 selected combinations not considered URM (e.g., White + Asian).
<sup>f</sup>PhD for highest degree completed includes 287 (78%) PhD and 28 (8%) MD/PhD, PharmD/PhD, or DVM/PhD; Other Doctoral includes 42 (12%) MD, DDS, or DMD, and 10 (3%) Other Doctorate.

Participants’ mean age was 36.9 ( ± 6.3) years. In terms of career stage, 271 participants (73%) were faculty and 96 (26%) were postdoctoral fellows (postdocs hereafter). Approximately two-thirds (66%) identified as female, and 46% identified as being from a URM background. A large proportion (86%) had a PhD as their highest degree (PhD only, MD/PhD, PharmD/PhD, or DVM/PhD). Approximately two-thirds were engaged in either laboratory-based (36%) or social behavioral (31%) research. The remainder were focused on clinical (22%) or epidemiological/data science research (10%).

The mean number of prior peer-reviewed publications was 19.6 ( ± 15.6). A majority of participants had submitted at least 1 prior training/career development application (60%) or research proposal (72%); 41% and 59% had received at least 1 prior training/career development grant or research grant, respectively. Additional details about participants’ grant application and funding history were previously reported [14].

### Rationale for examining faculty and postdocs outcomes separately

Although we present results for all participants, we focus on interpreting results for faculty and postdocs separately for two reasons: First, the interaction tests between intervention and career stage (faculty versus postdoc) on outcomes were not statistically significant; however, there was a trend for the structured (versus unstructured) effect to behave differently by career stage, particularly for K awards and success rates (*P* < .15). Second, we had an a-priori rationale: participants’ fundamental differences and the differential relevance of specific outcomes (eg, postdocs are not eligible to apply for R01 grants, whereas both faculty and postdocs are eligible to apply for K grants).

### General statistical considerations

Table 2 provides descriptive statistics (percentages) for the trial’s primary outcomes of submissions, awards, and success for any type of national-level grant application. Statistical differences in the experimental conditions were determined by the standard 2x2 factorial analysis. In this approach, logistic regression is used to test the effects of factor 1 (coaching dose: extended versus regular), factor 2 (SA engagement mode: structured versus unstructured), and their interaction on our primary outcomes (Table 3).

**Table 2.** Descriptive Percentages for Primary Outcomes, Defined as Any National-Level Grant Application and Award^a^, Overall and by Study Arm, for 367 US-Based Early-Career Faculty And Postdoctoral Fellows Enrolled in 2000-2022 Grant Coaching Intervention Study.

| Outcome Variable | Total Sample<br>(n=367) | Regular Dose<br>Unstructured<br>(n=90) | Regular Dose<br>Structured<br>(n=93) | Extended<br>Dose<br>Unstructured<br>(n=92) | Extended<br>Dose<br>Structured<br>(n=92) |
| --- | --- | --- | --- | --- | --- |
| **Self-Reported Submissions** |  |  |  |  |  |
| All Participants | 254/367 (69%) | 59/90 (66%) | 64/93 (69%) | 62/92 (67%) | 69/92 (75%) |
| Faculty | 198/271 (73%) | 46/68 (68%) | 51/71 (72%) | 48/66 (73%) | 53/66 (80%) |
| Postdocs | 56/96 (58%) | 13/22 (59%) | 13/22 (59%) | 14/26 (54%) | 16/26 (62%) |
| **Verified Awards** |  |  |  |  |  |
| All Participants | 116/367 (32%) | 22/90 (24%) | 29/93 (31%) | 25/92 (27%) | 40/92 (43%) |
| Faculty | 95/271 (35%) | 17/68 (25%) | 24/71 (34%) | 20/66 (30%) | 34/66 (52%) |
| Postdocs | 21/96 (22%) | 5/22 (23%) | 5/22 (23%) | 5/26 (19%) | 6/26 (23%) |
| **Success<sup>b</sup>**<br>**(Awardees / Submitters)** |  |  |  |  |  |
| All Participants | 116/254 (46%) | 22/59 (37%) | 29/64 (45%) | 25/62 (40%) | 40/69 (58%) |
| Faculty | 95/198 (48%) | 17/46 (37%) | 24/51 (47%) | 20/48 (42%) | 34/53 (64%) |
| Postdocs | 21/56 (38%) | 5/13 (38%) | 5/13 (38%) | 5/14 (36%) | 6/16 (38%) |

<sup>a</sup>National-level excludes proposals submitted to and awards funded by sources that do not convene national peer review panels, such as universities, local and state agencies, local and regional foundations, and health systems.
<sup>b</sup>Success is defined as any national-level award among those with any national-level submission.

**Table 3.**
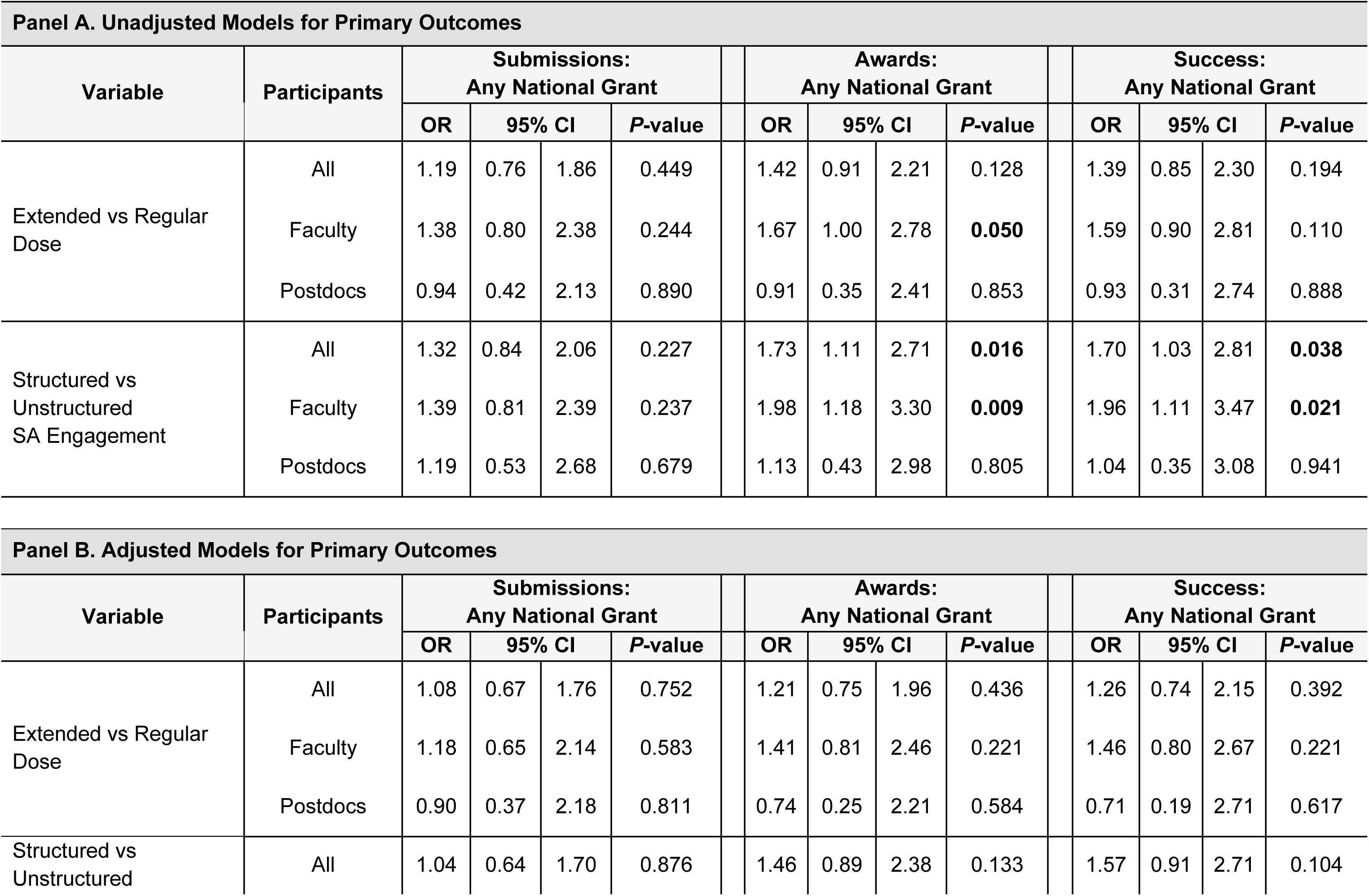
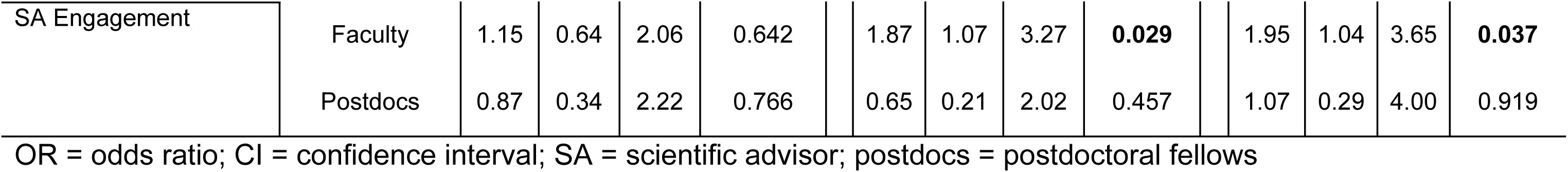
Primary Outcomes, Defined as National-Level Grant Application, Award, and Success Rate.

The following example illustrates our analytic approach. The data for faculty in Table 2 indicate that the rate of being awarded any national grant was higher in the extended dose structured arm (52%) than in the other 3 arms (25% to 34%). However, the 2x2 factorial analyses are needed to appropriately dissect the experimental effects. For example, if the combination of structured SA engagement and extended dose was statistically more effective than the other 3 arms, we would observe a significant interaction effect; alternatively, if the statistical driver of intervention impact is structured versus unstructured SA engagement, regardless of whether the dose was extended, we would observe a significant main effect of structured SA engagement and a non-significant interaction.

### National grant applications: submissions

Among participants, 254 (69%) submitted at least 1 national proposal by the 24-month follow-up timepoint (Table 2). Submission rates ranged from 66% to 75% across study arms and were higher for faculty than for postdocs (73% versus 58%). Descriptive statistics for secondary outcomes (specific grant mechanisms) are provided in S2-S4 Tables.

In the unadjusted and adjusted logistic regression models performed separately for faculty and for postdocs, submission rates for the primary outcome of any national grant application did not differ significantly by the 2 experimental factors (Table 3).

Differences were also not found in submission rates for major secondary outcomes that had sufficient prevalence to analyze in models (ie, K, R01 or R01-like) (Table 4).

**Table 4.**
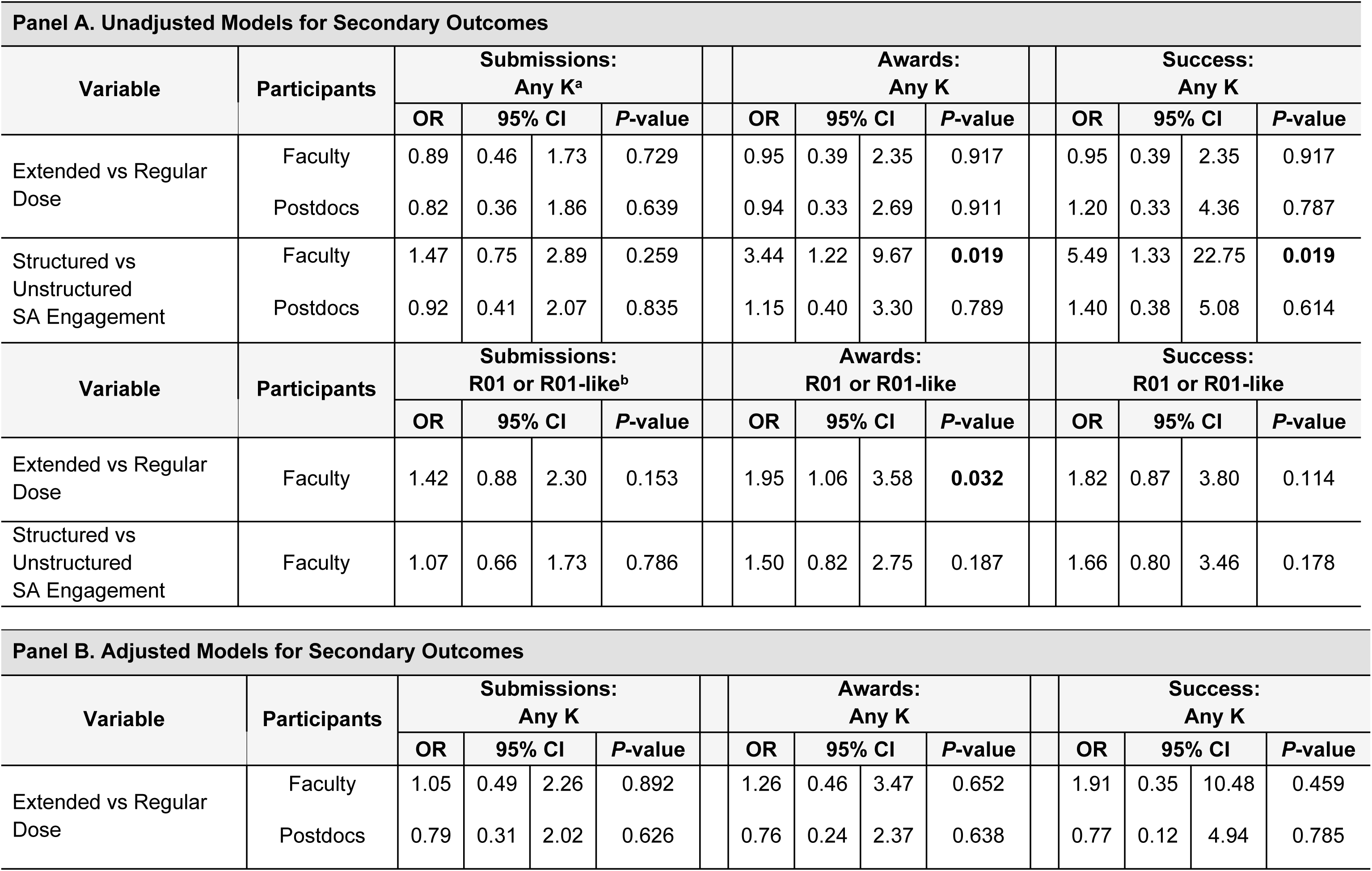
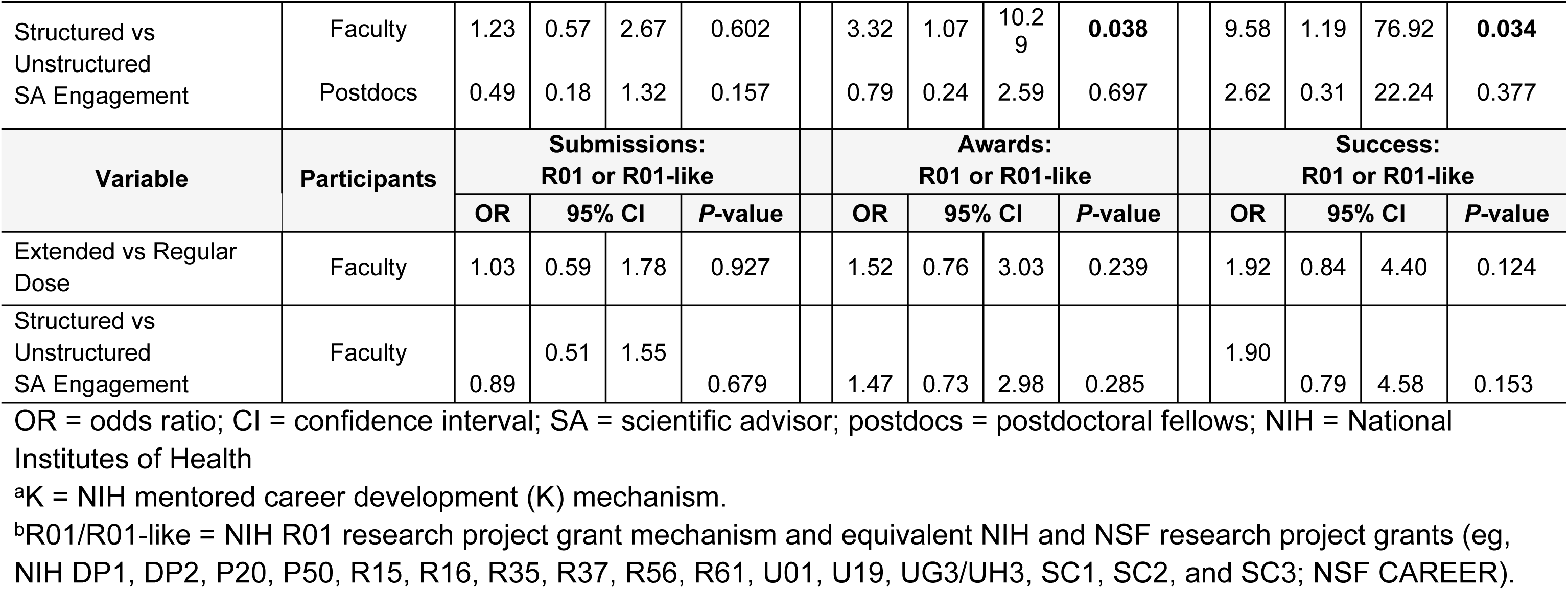
Major Secondary Outcomes, Defined as Submission, Award, and Success Rates for Specific Categories of National-Level Grants.

### National grant applications: awards

Among participants, 116 (32%) were awarded at least 1 national grant at 30-month follow-up (Table 2). Award rates ranged from 24% to 43% across study arms. For faculty, award rates across study arms ranged from 25% to 52%. In comparison, award rates were lower for postdocs, ranging from 19% to 23%. Twenty-six participants received more than 1 award; S1 Table provides a breakdown of the 147 verified grants that were awarded to 116 participants by funding agency and, for NIH awards, mechanism.

In the faculty models, participants in the structured (versus unstructured) SA engagement arms were more likely to have been awarded at least 1 national grant, with an effect size that was similar regardless of whether covariates were adjusted (unadjusted OR = 1.98, *P* = .009; adjusted OR = 1.87, *P* = .029; Table 3). This finding was driven by higher rates of at least 1 K award (unadjusted OR = 3.44, *P* = .019; adjusted OR = 3.32, *P* = .038; Table 4).

Differences in awards for faculty were also observed for intervention dosage in the unadjusted models. The odds of being awarded any national grant were higher for faculty in the extended (versus regular) coaching duration arms (unadjusted OR = 1.67, *P* = .050; Table 3), and nearly twice as high for the outcome of any R01 or R01-like award (unadjusted OR = 1.95, *P* = .032, Table 4). The odds ratios were still positive, although not significant, after adjusting for other covariates for any award (adjusted OR = 1.41, *P* = .221; Table 3) and for any R01 or R01-like award (adjusted OR = 1.52, *P* = .239; Table 4).

Both structured SA engagement and extended dose had a consistently positive moderate-to-large effect size (OR range 1.47 to 1.95) on faculty’s receipt of R01 or R01- like awards, even when not statistically significant, regardless of whether adjusting for covariates (Table 4).

In the postdoc models, there were no significant (unadjusted or adjusted) intervention effects on the primary outcome of any national award (Table 3) or any K award (Table 4).

### National grant applications: success rate

Lastly, we examined participants’ success rates, defined as awards received by those who submitted. Among the 198 faculty and 56 postdocs who submitted at least 1 national grant proposal, faculty success rates ranged from 37% to 64% across study arms, whereas postdocs had a consistent success rate of 38%, regardless of arm (Table 2).

In the faculty models, structured SA engagement, but not extended dose, doubled the success odds for those who submitted any type of national proposal (unadjusted OR = 1.96, *P* = .021; adjusted OR = 1.95, *P* = .037; Table 3). This was driven by the positive impact of structured SA engagement on faculty success in obtaining K awards (unadjusted OR = 5.49, *P* = .019; adjusted OR = 9.58, *P* = .034; Table 4). The effect size for both structured SA engagement and extended dose indicated a positive impact on R01 or R01-like success for faculty, regardless of whether adjusting for covariates (OR range 1.66 to 1.92; Table 4).

### Participants’ self-reported encounters with coaches during extended dose

Contrary to our hypothesis, the benefit of extended coaching on outcomes was not statistically significant. To explore this, we examined data from a feedback survey that was administered 24 months after the start of each cohort. Participants in the extended dose arms were asked, “During the Extended Dose period, what specific support did your coach provide?” to ascertain how many participants used their coaches and for what purpose. Specific response options and frequencies are provided in S5 Table. Response rates for this survey were low (50% for faculty, 36.5% for postdocs). Among these respondents, coach usage of any kind during the extended period was 79% for faculty and 63% for postdocs. Among faculty, usage was higher for those in the structured (versus unstructured) SA engagement arms.

### Multivariable associations between covariates and outcomes

It is common in randomized trials to explore the associations between baseline characteristics and outcomes, in addition to the causal impact of the experimental factors, to gain insights on potential catalysts for success that could be used to tailor future interventions. Although results for covariates are shown for the all-participant models (S6 Table), we focus on the results for faculty and postdocs separately (S7 and S8 Tables).

Among faculty participants, those engaged in laboratory-based research were significantly more likely to submit and be awarded R01 or R01-like grants, and less likely to submit and be awarded K grants, compared to those engaged in clinical research. Faculty who had submitted a training application prior to entering the study had higher odds of any submission and any R01 or R01-like submission, award, and success. This association was not observed for K submissions, awards, and success. Younger faculty were more likely to submit a K application, but older faculty were more likely to experience K success.

Among postdocs, prior training application submission was associated with higher odds of any K submission, and having any prior research application submission was associated with higher odds of obtaining any award.

### Intervention effect modifiers

Using only the final multivariable models in which experimental effects were adjusted for covariates, we examined intervention effect modification by testing the two-way interaction between each experimental factor and each covariate, adjusted for the other experimental factor and the other covariates. The covariates we examined as moderators included age, gender, URM status, career stage, research focus, prior publications, and previous submissions (training and research applications). S9 Table shows the significant interactions, including the overall interaction *P*-value as well as the intervention effects separately for different values of moderators (ie, simple effects). Although the all-participant results are shown in Panel A, we focus on the results for faculty and postdocs separately.

Among faculty (Panel B), the experimental factor of structured SA engagement had a significant interaction with the URM moderator on the outcome of any R01 or R01-like award (*P* = .030). The simple effects analysis, which provides the effect of the experimental factor separately for moderator categories, shows that structured SA engagement was approximately 4 times more effective than unstructured SA engagement for obtaining an R01 or R01-like award among faculty identifying as URM (OR = 4.32, *P* = .021). In contrast, structured SA engagement had no impact among non-URM faculty (OR = 0.81, *P* = .636). Likewise, structured SA engagement significantly improved success in obtaining any national grant (among those who submitted) for URM faculty (OR = 4.72, *P* = .003) but not for non-URM faculty (OR = 1.12, *P* = .779).

The other experimental factor, extended dose, had significant interactions with the previous research submissions moderator. Among faculty who had previously submitted a research proposal, extended dose coaching was 2 to 3 times more effective than regular dose for several outcomes: being awarded any R01 or R01-like grant (OR = 2.13, non-significant trend, *P* = .054) or achieving success on any R01 or R01-like grant (OR = 3.13, *P* = .017) or any national grant (OR = 2.08, *P* = .032). In contrast, faculty with no previous research submissions experienced no significant impact from extended coaching.

For postdocs (Panel C), there were 4 significant interactions between the structured SA engagement factor and previous submissions (either training or research proposals) on the outcomes of any submissions, any awards, and specifically K awards (*P*-values range .009 to .046 for the interaction tests). Those simple effects revealed that structured SA engagement was 2 to nearly 4 times more effective than unstructured SA engagement among postdocs who had no prior submissions (OR range 2.37 to 3.71, although simple effect *P*-values were not significant, ranging from .102 to .253). The opposite association was seen in postdocs with prior submissions, where structured SA engagement had a lower magnitude of impact than unstructured SA engagement (OR range .16 to .28; simple effect *P*-value was significant only for any submission outcome, *P* = 0.041).

## Discussion

Overall, the study’s results support the value of cross-institutional grant writing coaching groups, while providing insight into how this intervention might be further tailored to enhance its impact. The two experimental factors examined in the trial (coaching dose and mode of SA engagement) exhibited different effects, depending on the outcome being assessed and on participant career stage (faculty or postdoc). Key findings are as follows:

1. Neither factor influenced the outcome of proposal submissions.
2. Faculty with extended access to coaching had higher odds of being awarded any national grant or any R01 or R01-like grant in unadjusted bivariable models. These differences were no longer statistically significant after adjustment for covariates, but they consistently trended in the expected direction. The benefit of extended coaching was significant for one subgroup of faculty: those with a history of having submitted 1 or more previous research proposals before experiencing the intervention.
3. For faculty, structured SA engagement was associated in both unadjusted and adjusted models with significantly higher odds of being awarded any national grant or any K grant, and with significantly higher success rates among submitters of these proposal types. URM faculty, in particular, benefited from structured SA engagement, evidenced by a significant structured intervention effect that was not seen among non-URM faculty for awards (specifically, R01 or R01-like) and success rates (any national).
4. Postdocs differed from faculty in that awards and success rates were comparable across intervention arms.
5. Regardless of study arm, success rates (among submitters) were favorable, exceeding national rates for NIH K-series and R01 applications (shown below).

We discuss these findings next, including their implications for future interventions.

### Impact of coaching dose and scientific advisor engagement on outcomes

Successful submission of a proposal is a critical first step towards grants acquisition and can be a daunting task for less experienced investigators. Submission rates in our study sample were high regardless of assigned study conditions: 73% of faculty and 58% of postdocs submitted at least 1 national-level proposal within 30 months of the study.

Finding high submission rates among participants in grant writing coaching groups is consistent with previous results from our team [10] and institutionally based programs not designed as research studies [15,16]. The current study is the first to provide evidence from a national sample engaged in a randomized trial. Submission rates in our study were likely boosted by a self-selected sample of participants already motivated to submit grant applications. We also enrolled participants based on their readiness to write. This means our submission results might not be generalizable to an “all comers” approach to enrollment in this kind of intervention. We did not observe significant differences in submission rates between study arms. This finding suggests that the core elements of our prototype intervention, offered in 5 months of group coaching, provided as much benefit as possible with respect to facilitating submissions.

In contrast to submissions, awards and success rates differed by study arm, but only among faculty participants (not postdocs). Both outcomes were highest in the extended dose + structured SA engagement arm, approximately double those of faculty in the regular dose + unstructured SA engagement arm (52% versus 25% for awards, 64% versus 37% for success rates). Our multivariable 2x2 factorial analyses suggest this difference was driven primarily by the effect of structured SA engagement, particularly for faculty developing K proposals, rather than by extended access to a coach.

We can speculate why faculty, and not postdocs, benefited from structured SA engagement, especially those writing mentored K applications. A reasonable assumption is that early-career faculty have less access to mentorship than postdocs, but a successful K application depends heavily on mentor input. Therefore, a structured protocol such as ours—one that proactively prompts mentors to engage with their faculty mentees at specific stages of proposal development—might help to encourage earlier, more frequent, and more productive interactions than might otherwise occur. Additionally, faculty writing K applications are at earlier stages of developing their research ideas than faculty writing R01 or similar research proposals. Thus, a structured protocol designed to solicit and align feedback from multiple readers (coach, SA, coaching group participants, mock reviewers) offers particular value to faculty preparing K proposals. By comparison, postdocs writing K awards, by definition, have mentors, as they are still in a dependent career stage.

One explanation for not seeing a consistent benefit of the extended dose effect is the uneven usage of coaches in months 6-18 by faculty and postdocs (79% and 63% of survey respondents, respectively, S5 Table). Aside from initial outreach by the coach at the start of the extended dose period, all interactions were participant-driven. It is plausible that many participants, particularly postdocs with well-established mentorship and individuals working on K applications, found it less valuable to continue engaging with a grant writing coach from outside their institution beyond the initial, 5-month grant writing period, preferring instead to refine their drafts (and any subsequent revisions) in close collaboration with their local mentors. In addition, participants’ decision to engage coaches beyond the regular dose may have depended on the timing of their initial or re- submission. Rates of coach usage were higher for faculty in the structured SA engagement arms than those in the unstructured arms (but not significantly so, *P* = .099). The explanation for this is not obvious, although the encounters between SAs and coaches during the regular dose period may have helped to reinforce the coach’s value, thereby encouraging extended usage.

### Simple effects analyses of intervention modifiers

Two major themes emerged from the moderator subgroup analyses for faculty. First, URM faculty benefited significantly more from structured SA engagement than non-URM faculty in terms of R01 or R01-like awards and success rates for any national proposal. We do not have a clear explanation for this finding, although previous studies have identified barriers for URM individuals obtaining access to mentors [11,17,18]. Second, extended coaching dose had a more positive impact (ie, greater R01 or R01- like awards and success) for faculty with prior submissions than those without. This suggests that faculty with prior submissions may have been further ahead in their readiness to engage with coaches than those without prior grant submissions. Ongoing assistance from coaches with interpreting summary statements and preparing resubmissions could have had important positive effects on outcomes for these investigators.

### Comparison of success rates to NIH data

Regardless of study arm, success rates for national applications among faculty in our trial were high, exceeding rates reported by the NIH for K-series and R01 and R01-like applications. For the period 2020 to 2024, the average success rate for new R01 and R01-like proposals was 17.6% for first-time investigators [19]. The composite success rate for faculty R01 and R01-like proposals across the 4 arms of our study was 43% (excluding NSF awards), a 2.5-fold difference. Over this same period, the average success rate of K01, K08, and K23 awards by NIH data was 36.5% compared to 51% for faculty in our study [20]. The difference is less substantial than that for R01 and R01- like awards. This could reflect the fact that those writing K awards, by definition, have mentors assisting them with the proposals; thus, we might expect less of an impact of the supplemental intervention. However, qualitative interviews with faculty writing K applications revealed substantial positive value for some (manuscript currently under review).

### Implications of study findings

Our results have several implications for the design and implementation of interventions to support the grant-seeking efforts of biomedical researchers:

1. *Demand is high for cross-institutional grant writing coaching groups.* We had to turn away many applicants who wanted to be part of the study, and we continue (in 2026) to receive inquiries about whether more groups will be offered. Large, high-resourced institutions may have the infrastructure and numbers of early-career investigators to enable local grant writing groups. Smaller, lower-resourced institutions may lack a critical mass of participants and mentors, creating a significant disadvantage for investigators at these sites. Our national model levels the playing field. Virtual coaching groups enable cross-institutional, ongoing (versus one-time) coaching, avoiding limitations such as the time and costs of travel.
2. *This national group coaching model demonstrates an especially high impact for faculty trying to reach the R01 level.* This is a big step coming off a K award or starting without one. Overall, the model is valuable for junior faculty who are at this make-or-break career juncture.
3. *Group facilitation by a highly skilled coach is critical, but the degree to which participants engage with the coach after the regular, immersion group phase is variable.* Having extended access to a coach is valuable to some participants, but not all. Thus, expectations for coach usage over an extended period should be modest. Our qualitative interviews with participants and coaches revealed that ongoing coach engagement was most likely when a strong connection developed during the group coaching period, and when the fields of the coach and participant aligned well–conditions that were not universal (manuscript currently under review).
4. *Some degree of structured integration of scientific advisors with the coaching process can be beneficial*. Per our findings, URM faculty and faculty preparing K awards are two groups that may derive the most benefit from this integration.

### Strengths and limitations

We successfully enrolled a large, diverse sample (by gender, URM status, and research focus) across numerous institutions to test variations of a grant writing coaching group intervention using a randomized trial design. The literature includes many single program evaluations [15,16,21,22], which are helpful but lack a robust study design intended to tease out what elements of intervention design are most effective and for what groups; our study is highly innovative in that it fills this gap.

Study limitations include the lack of a true control group (ie, usual care: no coaching). It was the team’s judgement that having a usual care study arm was not ethical, given substantial data on the efficacy of the general coaching approach. Therefore, having 4 arms with treatments may have limited the ability to find between-arm differences, which is the rationale for using historical national NIH data to show overall improvement over usual care. Another limitation is that although arms were randomized, it is possible that arms were unbalanced at baseline on unmeasured variables, which, if adjusted as covariates, could have reduced residual variation and therefore either strengthened or weakened the between-arm differences. In addition, including faculty and postdocs in the same study limited the sample size and power for analyses of each separately.

## Conclusion

There is strong support for the core prototype model used in the study. A high number of participants submitted national-level grant applications, and those who submitted did well compared to NIH funding rates. Our trial’s 2x2 factorial design, with its national scope and inclusion of both faculty and postdocs, yielded insight into how the standard intervention might be optimized.

## Acknowledgements

The authors thank biostatistician Marlene J Egger, PhD, for her significant contributions to this study until her retirement in 2022; NIH staff Jessica M Faupel-Badger, PhD, for consulting on this study; David Guise for data management support, and the grant writing coaches who played essential roles in the implementation of this study.

## Supporting information captions

S1 Table. Awards by Source (and Mechanism for NIH) for Faculty and Postdocs. NIH = National Institutes of Health. NSF = National Science Foundation.

**S2 Table. Descriptive Statistics for Submission Outcomes (Primary and Secondary).** Data are presented as numbers with percentages (%) in parentheses.

**S3 Table. S3 Table. Descriptive Statistics for Award Outcomes (Primary and Secondary).** Data are presented as numbers with percentages (%) in parentheses.

**S4 Table. Descriptive Statistics for Success Outcomes (Primary and Secondary).** Data are presented as numbers with percentages (%) in parentheses. Success = any national-level award among those with any national-level submission of corresponding grant type.

**S5 Table. Extended Dose Responders to 24-Month Coaching Survey.**

**S6 Table. All Participants: Multivariable Logistic Regression Models.** Any success = any national-level award among those with any national-level submission of corresponding grant type. OR = adjusted odds ratio. NE = not estimable for OR (95% CI) due to small sample size. URM = underrepresented in medicine. Epi/DS = epidemiology/data science research focus. Soc/Beh = social/behavioral science research focus. ^a^*P*-value is from logistic regression’s two-sided Wald test.

**S7 Table. Faculty: Multivariable Logistic Regression Models.** Any success = any national-level award among those with any national-level submission of corresponding grant type. OR = adjusted odds ratio. NE = not estimable for OR (95% CI) due to small sample size. Uns = unstructured. Reg = regular. Early = early career. URM = underrepresented in medicine. Epi/DS = epidemiology/data science research focus. Soc/Beh = social/behavioral science research focus. Train = training. Res = research. Subs = submissions. Pubs = publications. ^a^*P*-value is from logistic regression’s two-sided Wald test.

**S8 Table. Postdocs: Multivariable Logistic Regression Models.** Any success = any award among those with any submission of the corresponding award type. URM = underrepresented in medicine. NE = not estimable for odds ratio (95%) due to small sample size. Epi/DS = epidemiology/data science research focus. Soc/Beh = social/behavioral science research focus. ^a^*P*-value is from two-sided Wald test from logistic regression.

**S9 Table. Simple Effect for Explaining Significant Interactions.** Inter = interaction. URM = underrepresented in medicine. NA = not available from SAS LOGISTIC SLICE procedure; refer to confidence interval. NE = not estimable. SA = scientific advisor.

